# Whole genome sequencing of dog specific assemblages C and D of *Giardia duodenalis* from single and pooled cysts indicates host associated genes

**DOI:** 10.1101/645119

**Authors:** F.N.J. Kooyman, J.A. Wagenaar, A. Zomer

## Abstract

*Giardia duodenalis* (Syn. *G. intestinalis* or *G. lamblia*) infects over 280 million people each year and numerous animals. *G. duodenalis* can be subdivided into 8 assemblages with different host specificity. Unculturable assemblages have so far resisted genome sequencing efforts. In this study we isolated single and pooled cysts of assemblage C and D from dog faeces by FACS and sequenced them using multiple displacement amplification and Illumina paired end sequencing. The genomes of assemblages C and D were compared with genomes of assemblages A and B from humans and assemblage E from ruminants and pigs. The genomes obtained from the pooled cysts and from the single cysts were considered complete (>99% marker genes observed) and the allelic sequence heterozygosity (ASH) of assemblage C and D was 0.89% and 0.74%, respectively. Higher than for assemblage B (> 0.43%) and much higher than for assemblages A and E (<0.01%). The flavohemoglobin and 4Fe-4S binding domain family gene involved in O_2_ and NO detoxification were only present in assemblages A, B and E. Cathepsin-B orthologs were found in all genomes. Six clades of cathepsin-B orthologs contained one gene of each genome, while in three clades not all assemblages were represented. We conclude that whole genome sequencing from a single Giardia cyst results in complete draft genomes making the genomes of unculturable Giardia assemblages accessible. Observed differences between the genomes of assemblage C and D on one hand and the assemblages A, B and E on the other hand are possibly associated with host specificity.

## Introduction

*Giardia duodenalis* (*syn. G. lamblia, G. intestinalis*) is a common intestinal parasite of mammals, including man, with diarrhoea being the most common symptom. More than 280 million annual cases of human infections are described (Lane and Lloyd 2002) and numerous cases in other mammals. In *G. duodenalis* 8 assemblages (A to H) have been described up till now, with various degree of host specificity (Sprong et al. 2009; Feng and Xiao 2011). Their genetic differences and host specificities has made some researchers suggest that some of the assemblages can be considered separated species (Franzén et al. 2009; Jerlström-Hultqvist et al. 2010a; Monis et al. 2003).

The assemblages A and B are primarily found in humans, but have occasionally been found in dogs as well. The assemblages specific for dogs are the assemblages C and D. In humans only 0.3% of infections consisted out of assemblage C or D (Sprong et al. 2009). Assemblage A can be divided into sub-assemblages AI, AII and AIII, of which AI occurs both in man and other mammals, like dogs, while AII occurs almost exclusively in humans (Sprong et al. 2009; Feng and Xiao 2011). Assemblage B is diverse with no clear sub-assemblages. Assemblage AI, AII and B are assumed to be zoonotic.

The genome of *G. duodenalis* is a compact genome with little non-coding regions and only 4 introns (Morrison et al. 2007). The parasite is supposed to be a clonal reproducing organism with 2 diploid, almost identical nuclei. Without exchange of DNA between the 2 nuclei, random mutation will result in increasingly sequence differences between the 2 nuclei and thus increasingly heterozygosity within the individual. However, the level of heterozygosity in *Giardia* was lower than expected (Morrison et al. 2007) and the limited exchange of DNA between the nuclei, called diplomixis by Poxleitner et al. (2008) can be the cause for this low level of heterozygosity. The allelic sequence heterozygosity (ASH) vary in the different assemblages from very low (< 0.0023%) in assemblage E to much higher (0.53%) in assemblage B (Franzén et al. 2009). The cause or the consequence of these differences are unknown.

Of the 8 assemblages of *G. duodenalis*, only 3 assemblages have their genome sequenced and published. Two genomes of assemblage A have been sequenced: one from assemblages AI, strain WB (Morrison et al. 2007) and one from assemblage AII, strain DH (Adam et al. 2013). From assemblage B, there are also 2 isolates sequenced, GS (Franzén et al. 2009) and GSB (Adam et al. 2013). Of the non-human assemblages, only assemblage E, strain P15 from a pig is sequenced (Jerlström-Hultqvist et al. 2010b). This assemblage is associated with artiodactyls (ruminants and pigs). All the genomes are comparable in size ranging from 10.7 to 12.0 Mb for the haploid genome and between 5,008 and 7,477 ORFs (Adam et al. 2013). Differences between the assemblages in genome size and number of ORFs can be a biological difference, however different sequencing methods and subsequent assembly and gene calling can add to this differences as well.

The mechanisms behind the host specificity of the assemblages is not known, nor why some infections are without symptoms, while in other cases severe diarrhoea develops upon infection. Proteins excreted by the trophozoites are likely to play a role in host specificity and immunity. Several proteins from the WB and GSB isolates excreted in the absence of host cells were identified by proteomics (Dubourg et al. 2018). The most abundant proteins in both isolates were variant-specific surface proteins (VSPs), cathepsins, high cysteine membrane proteins and tenacins. Differences in proteolytic activity and specificity of the cathepsin paralogs and orthologs of different isolates/assemblages have been described (Bhargava et al. 2015; Lui et al. 2018). So far, nothing is known about the cathepsins of the assemblages of the dog.

The whole genome sequences and the proteomic data so far were all derived from cultured *G. duodenalis* trophozoites. Bases on whole genome sequences, assemblage specific proteins could be identified by proteomics and characterised. Assemblages from the dog (C and D), the cat (F), the rat (G) and from the pinnipeds (H) have resisted culturing so far. Therefore, other methods for obtaining DNA of sufficient quality and quantity for whole genome sequencing are needed. Purification of cysts from a clinical isolate by immunomagnetic beads is an option. This method has resulted in sufficient pure *Giardia* DNA for whole genome sequencing (Hanevik et al. 2015), but (unnoticed) mixed-infections will give spurious results. This can be circumvented by starting with the isolation of a single cyst followed by high quality amplification of the small amount of genomic DNA. Troell et al. (2016) was able to isolate individual oocysts of *Cryptosporidium parvum* prior to whole genome amplification and sequencing. Single cell coverage was on average 81% and by combining 10 individual cells the whole genome was covered.

The aim of the study was to obtain the whole genome sequences of the dog specific assemblages C and D of *G. duodenalis* and to perform a comparative analysis with the genomes from the already sequenced assemblages. Hereto, the DNA from single cysts and pooled cysts (control for completeness) was amplified with multiple displacement amplification (MDA). Whole genome sequencing was performed on the resulting MDA product and the genomes were assembled and annotated. Finally, the genomes from assemblage C and D obtained from single and pooled cyst were compared with each other and with the already known genomes from the assemblages A, B and E.

## Material and Methods

### Dogs and parasites

*Giardia* positive faecal samples from 9 dogs were obtained from the Veterinary Microbiological Diagnostic Centre (VMDC, Faculty of Veterinary Medicine, Utrecht University, The Netherlands).

### Isolation of cysts

Faeces were collected and stored at 4◻C for no more than 1 week, before the *Giardia* cysts were isolated. Isolation of the cysts was done by suspending 1 to 5 g of faces in approximately 25 ml H_2_O and filtered over an open filter chamber with mesh size of approximately 100 μm (Beldico, France). The filtrate was centrifuged for 2 min at 2,000 g and the pellet was resuspended in 50 ml H_2_O. Centrifugation and resuspending were repeated until the supernatant was clear. Finally, the pellet was resuspended in 25 ml H_2_O. Ten ml sucrose solution (70 g sucrose in 100 ml H_2_O) was applied at the bottom of the tube without mixing with the cysts-suspension. The tube was centrifuged without break for 15 min at 2,000 g. The resulting interphase, containing the majority of the *Giardia* cysts was harvest, washed again with H_2_O as described and resuspended in 1 ml of PBS. Subsequently, the cysts were counted in a modified Fuchs-Rosenthal counting chamber. Samples from each isolate were preserved in 70% EtOH at 4◻C as well as frozen in 33 μl samples at −20 ◻C.

### Isolation of DNA from cysts

DNA isolation from sucrose purified cysts was performed with QIAamp Fast DNA Stool Mini kit (Qiagen) as described for dog faeces (Uiterwijk et al. 2018) with the following modification: Only 33 μl frozen purified cysts was used as starting material, instead of 0.2 g of faeces and only 167 μl of inhibition/extraction buffer was used.

### Determination of assemblage by multilocus genotyping (MLG)

The *bg* and *gdh* loci were amplified by nested PCR. All amplifications were performed with 2.5 μl template and Dreamtag polymerase (Thermoscientific) supplemented with 0.5 mg/ml BSA (Sigma, catalog nr. B287). Nested PCR on the *gdh* locus was performed with primers described by Cacciò et al. (2008). Conditions for primary amplification: 35 cycles (94 ◻C for 30 s, 57.5 ◻C for 30 s and 72 ◻C for 60 s). The undiluted primary product (2.5 μl) was used as template for the secondary amplification. Conditions for secondary amplification: 35 cycles (95 ◻C for 30 s, 60 ◻C for 30 s and 72 ◻C for 60 s). Each amplification started with a denaturation step of 95 ◻C for 3 min and ended with a final extension step of 72 ◻C for 7 min. PCR on *bg* locus was performed with primers as described by Tseng et al. (2014). Conditions for primary amplifications were: 35 cycles (95 ◻C for 30 s, 65 ◻C for 30 s, 72 ◻C for 60 s). Primary products were 10x diluted before use as template in secondary amplification. Conditions for secondary amplification were: 35 cycles (95 ◻C for 30 s, 50 ◻C for 30 s, 72 ◻C for 60 s). Each amplification started with a denaturation step of 95 ◻C for 3 min and ended with a final extension step of 72 ◻C for 7 min. The amplicons were treated with EXO-SAP (affimetrix) and Sanger sequencing was performed at BaseClear (Leiden, The Netherlands). The assemblage was determined by aligning the sequences of the amplicons with reference sequences of assemblages A to G (Sprong et al., 2009) with the ClustalW module in DNASTAR Lasergene 14.0 ©.

### Labelling of cysts and FACS

One ml of EtOH preserved cysts was added to 100 μl of zirconium beads (diameter 0.5 mm). The cysts were washed twice with 1 ml PBS by centrifugation (2 min, 2,000 g) and aspirated without disturbing the beads. PBS (170 μl) and 2 drops of detection reagents (Merifluor C/G), containing FITC labelled mAb against *Giardia* and *Crytosporidium,* were added to the beads-cysts suspension. The suspension was mixed gently end over end, stained for 30 min at room temperature and washed again twice as described. Finally, 200 μl PBS was added to the mixture, mixed gently and the cysts were transferred in 200 μl (without centrifugation), while taken care not to transfer the beads. The cyst suspension was poured through a 100 μm cell strainer (Greiner bio-one) and checked for stained *Giardia* cysts and the absence of *Cryptosporidium* with fluorescence microscopy. Cysts were counted in modified Fuchs-Rosenthal counting chamber and brought to an concentration of 5 cysts per μl PBS.

### Sorting of Giardia cysts

First, *Giardia* cysts were identified and gated based on the FSC and SSC profile (Figure S1). Within this gate, doublets were eliminated based on FSC and pulse width parameters. Finally only the fluorescently labeled particles were sorted based on FSC and FITC-fluorescence parameters. Single cysts and pools of 10 cysts were collected in wells containing 2 μl PBS.

### Multiple Displacement Amplification (MDA)

Individual single *Giardia* cysts or pools of 10 cysts isolated by FACS were subjected to MDA with the REPLI-g single cell kit (Qiagen). MDA was performed according to the manual, except that all volumes were halve of the prescribed volumes. Confirmation of MDA amplification of the right assemblage was achieved by PCR using only the first primer pairs of the *gdh* and *bg* loci on the 100x diluted MDA product and subsequent sequencing of the amplicons. Sequences were aligned with the reference sequences (Sprong et al. 2009). Control for contamination by small amounts of assemblage C in MDA products of assemblage D or vice versa, was performed by RFLP on the amplified *bg* fragment. The *bg* fragment of assemblage D (AY545647) and assemblage C (AY545646) differ in the number of restriction sites for XhoI. Pooling of cells for MDA reactions (Ellegaard et al. 2013) as well as pooling of MDA products for sequencing reactions (Lasken, 2007) can increase the completeness of the genome. Therefore, 4 MDA products, each obtained from 10 cysts of the same assemblage from the same dog were, when proven to be without contamination, pooled into one vial. This results in pooled MDA products of 4×10 cysts. MDA products obtained from individual cysts were kept separately. All the MDA products were subsequently purified on QIAampDNA mini kit and eluted with AE buffer. dsDNA contents was determined with Qubit and all samples were diluted to 200 ηg dsDNA in 50 μl AE buffer.

### Whole genome sequencing and genome analysis

WGS was performed on Illumina MiSeq platforms (Illumina, USA) using 2 × 250-bp reads with approximately 230 fold coverage per genome. 85% of the reads could be aligned to the nonredundant (nr) protein database using DIAMOND (Buchfink et al. 2015) and classified using MEGAN (Huson et al. 2016) to evaluate contamination with bacterial DNA. Approximately 0.25% of the reads were classified as bacterial and 99.75% as eukaryotic of which 99.98% was assigned to Hexamitidae (Figure S2). Genomes were assembled with SPAdes v3.10.1 (Bankevich et al. 2012) using default settings. Contigs with a size <200 bp and a coverage lower than 10 were removed. All contigs were aligned with the nr protein database. Contigs classified as bacterial by MEGAN were removed (approx. 0.7% of the contigs). Genes were identified using Prodigal v2.6. of the sequenced Giardia genomes and protein annotations were collected from GiardiaDB (https://giardiadb.org/giardiadb/) and all versus all BLAST was performed for all predicted proteins of the genomes and *Spironucleus salmonicida* as outgroup at an E-value cutoff of 1E-6. To determine the orthologous relationships of all proteins, the BLAST output was parsed by Orthagogue (Ekseth et al. 2014). Proteins were considered for orthology clustering if the proteins had at least 20% identity and at least 20% overlap. To determine the orthologous groups (OGs), Markov clustering (MCL) was performed using MCL-edge (Enright et al. 2002). Proteins were aligned with each other within their respective OGs using MUSCLE (Edgar 2004) and the gene names and functions from the reference genomes were added. Completeness of the genomes was assessed by calculating the recovery of the single copy genes, present in all the genomes of the assemblages A, B and E (n= 2716).

A protein super alignment of 534,584 positions was created by concatenating the aligned proteins if they were present in all assemblages and the outgroup *S. salmonicida*. Phylogenetic dendrograms were created using IQTREE using the automated model selection method (-m TEST), 1000 ultrafast bootstrap cycles (-bb 1000) and visualized using MEGA 6.6.

Venn diagrams were constructed by on line webtool from University of Ghent, Belgium (http://bioinformatics.psb.ugent.be/webtools/Venn). The alignment and phylogeny of the cathepsin-B family was performed in MEGA 6.6.

## Results

### Dogs and parasites

Sucrose purified cysts from nine *G. duodenalis* infected dogs were characterized by MLG. Out of the 9 dogs, 3 dogs turned out to be infected with *G. duodenalis* assemblage D, 3 dogs were infected with assemblage C and 3 dogs were infected with a mixture of assemblage C and D (not shown). Isolated cysts from 3 dogs were selected for FACS and MDA: Dog 8 (galgo español, female, 7 months old) was infected with assemblage C and was suffering from mild diarrhoea, dog 5 (French bulldog, female, 6 months old) was infected with assemblage D and was suffering from watery diarrhoea and dog 1(bull terrier, female, 6 months old) was infected with a mixture of assemblage C and D and was suffering from chronicle hemorrhagic diarrhoea.

### FACS and MDA

The individually sorted single cysts from dog 1 (assemblage C or D) and the cysts sorted in pools of 10 from dog 5 (assemblage D) and dog 8 (assemblage C), were subjected to MDA amplification. From all MDA products from pooled cysts the amplification of the single copy loci *bg* and *gdh* by PCR was successful (not shown). Seven out of the 8 single cysts from dog 1 also yielded MDA products suitable for amplification of both single copy loci. From these 7 cysts that yielded a PCR product, 4 cysts were of assemblage C and 3 were of assemblage D (not shown). Because these single cysts were obtained from the same dog with a mixed infection, contamination of the single cysts with DNA of the other assemblage can be a possibility. However, PCR fragments amplified from MDA products of single cysts showed no “double peaks” at the positions were assemblage C and D differ. Furthermore, in none of the MDA products contamination by DNA of the other assemblage could be demonstrated by PCR/RFLP (not shown).

### Characteristics of the genomes

The characteristics of the genomes are given in table 1 and the presence absence score is given in table S1. The sequencing of the genomes of assemblage C resulted in 3,388 to 3,917 contigs with a coverage of 206 to 235. The genome was estimated to be between 11.5 and 12.1 Mbp. The sequencing of the genomes of assemblage D resulted in 2,885 to 3,489 contigs with a coverage of 231 to 267. The genome was estimated to be between 11.4 and 11.5 Mbp. All genomes contained about 3900 OGs.

**tabel 1.** Characteristics of *G. duodenalis* genomes.

| Genome | Assemblage | nr of reads | total bases | coverage | Genome size (bp) | nr of contigs | nr of genes | nr OGs | Completeness (%) | ASH (%) | Reference |
| --- | --- | --- | --- | --- | --- | --- | --- | --- | --- | --- | --- |
| WB | AI | ± 200,000 | 1,41E+08 | 11 | 12,827,416 | 211 | 6,027 | 4,199 | ND | <0.01 | Morrison et al. 2007 |
| DH | AII | 4,996,200 | 1,33E+09 | 124 | 10,703,894 | 239 | 5,147 | 3,505 | ND | 0.037 | Adam et al. 2013 |
| GS | B | 808,181 | 1,76E+08 | 16 | 11,001,532 | 2,931 | 4,471 | 3,469 | ND | 0.53 | Franzen et al. 2009 |
| GSB | B | 3,019,027 | 6,37E+08 | 53 | 12,009,633 | 543 | 6,094 | 3,803 | ND | 0.43 | Adam et al. 2013 |
| P15 | E | 1,639,140 | 5,42E+08 | 47 | 11,522,052 | 820 | 5,068 | 3,609 | ND | 0.0023 | Jerlstrom-Hultqvist et al. 2010b |
| SSAL |  | 2,390,565 | 5,18E+08 | 40 | 12,954,588 | 233 | 6,288 | 3,997 | ND | ND | *Xu et al. 2014* |
| cyst 1 | C | 7,903,382 | 2,38E+09 | 206 | 11,557,310 | 3,388 | 5,550 | 3,993 | 99.3 | 0.85 | *This paper* |
| cyst 3 | C | 9,026,378 | 2,72E+09 | 235 | 11,550,696 | 3,651 | 5,391 | 3,961 | 99.0 | 0.88 | *This paper* |
| pool 8 | C | 8,821,718 | 2,66E+09 | 220 | 12,061,379 | 3,917 | 5,538 | 3,986 | 99.2 | 0.94 | *This paper* |
| cyst 2 | D | 8,714,436 | 2,62E+09 | 231 | 11,374,926 | 3,269 | 5,350 | 3,923 | 99.1 | 0.80 | *This paper* |
| cyst 4 | D | 10,000,252 | 3,01E+09 | 267 | 11,268,649 | 2,885 | 5,460 | 3,974 | 98.6 | 0.63 | *This paper* |
| pool 5 | D | 9,496,842 | 2,86E+09 | 249 | 11,499,674 | 3,489 | 5,549 | 4,016 | 99.0 | 0.78 | *This paper* |

nd= not determined

### Completeness of genomes

Single copy genes are defined here as an OG with exactly one member in all 5 previously sequenced genomes WB, DH, GS, GSB and P15. There were in total 2716 single copy genes and of these genes, 99.0% −99.3% and 98.6% −99.1% were present in the genomes of assemblages C and D, respectively (Table 1). No differences in completeness between genomes derived from single cysts or from 40 pooled cysts were found.

### Shared and unique OGs in assemblages C and D

The unique and shared OGs from genomes obtained from individual and pooled cysts were identified for assemblages C and D. In the 3 isolates there were 4266 and 4203 OGs identified in assemblage C and D, respectively (Figure 1). The vast majority of OGs (88%-89%) were shared by all 3 isolates of the same assemblage and 92.9%-93.3% of the OGs were shared by at least 2 isolates of the same assemblage. The number of unique OGs were slightly higher in the genomes obtained from the pooled cyst compared to the genomes from the single cysts.

**Fig. 1.**
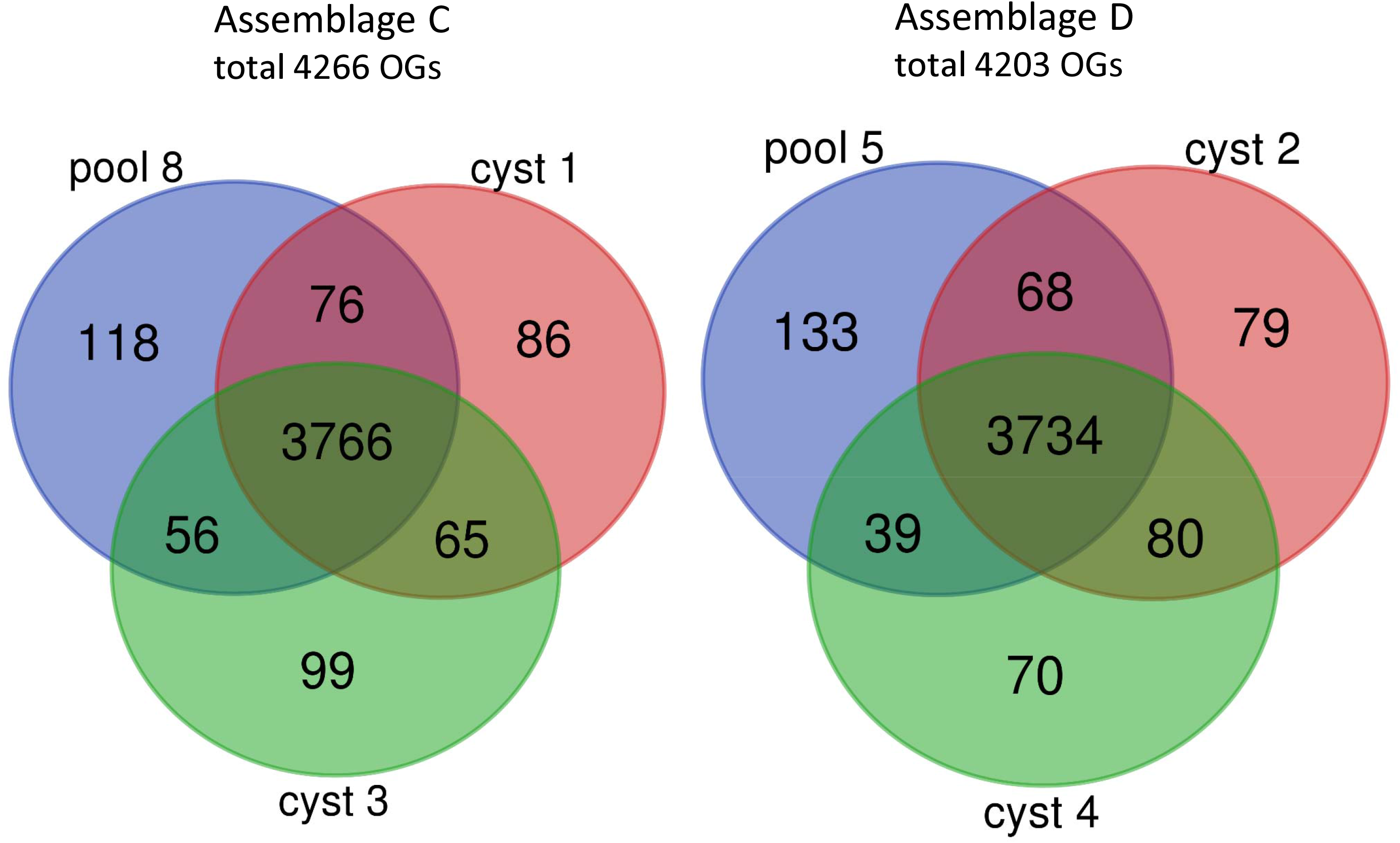
Unique and shared OGs in assemblage C and D. Distribution of the OGs of the 3 genomes within assemblage C (pool8, cyst 1 and cyst 3) and within assemblage D (pool 5, cyst 2 and cyst 4) shown in a Venn diagram. The genomes from cyst 1, 2, 3 and 4 were obtained from MDA products of single cysts. The genomes from pool 8 and pool 5 were obtained from pooled MDA products of 40 cysts per assemblage.

The proteins in all 6 genomes from assemblage C and D were compared with each other (Figure S3). There were in total 4859 OGs of which 3485 OGs (71.7%) were shared between all genomes. Furthermore, 243 OGs (5.0%) and 194 OGs (4.0%) were specific for assemblage C and D, respectively. Although the sequencing of the PCR products and the RFLP of the MDA product did not reveal cross contamination, the genomes were also inspected for this potentially contamination. OGs present in all the genomes from the individual cysts, but absent in one of the genomes from the pooled cysts is indicative for contamination in the individual cysts. Only 11 (0.23%) and 10 (0.21%) OGs were present in all the genomes from the individual cysts, but absent in genomes from pooled cysts of assemblage C and assemblage D, respectively. In genomes of single cyst 1 and cyst 3 from assemblage C there were respectively, 9 and 21 OGs absent (mean 0.31%) and from genomes of single cyst 2 and 4 from assemblage D there were respectively, 2 and 19 OGs (mean 0.22%) absent. This all indicates that contamination of the genome of one assemblage with that of the other assemblage is, if any, very limited. A direct comparison of unique and shared OGs between the genomes of the single cysts and pooled cysts is also given in figure S4.

### Allelic sequence heterozygosity (ASH)

The allelic sequence heterozygosity (ASH) defined by an alternative base calling higher than 15% and a 10-fold coverage or higher (Adam et al. 2013), resulted in a mean ASH for the 3 genomes of 0.89% and 0.74% for assemblage C and D, respectively (table 1). The heterozygosity was similar for the genomes from single cysts and pooled cysts, but was very much depending on the threshold for alternative base-calling (Figure 2A). The ASH for assemblage C and D was slightly higher than for GS (0.53%) and much higher than for WB (<0.01%) and P15 (0.0023%) (Franzén et al. 2009; Adam et al. 2013). For 5 out of the 6 genomes, the number of SNPs decreased linear with the percentage of alternative base calling (Figure 2B). Only for the genome from pool 8 (assemblage C) the number of SNPs peaked by an alternative base calling of around 25%, suggesting the presence of 4 haploid genomes in this sample.

**Fig. 2.**
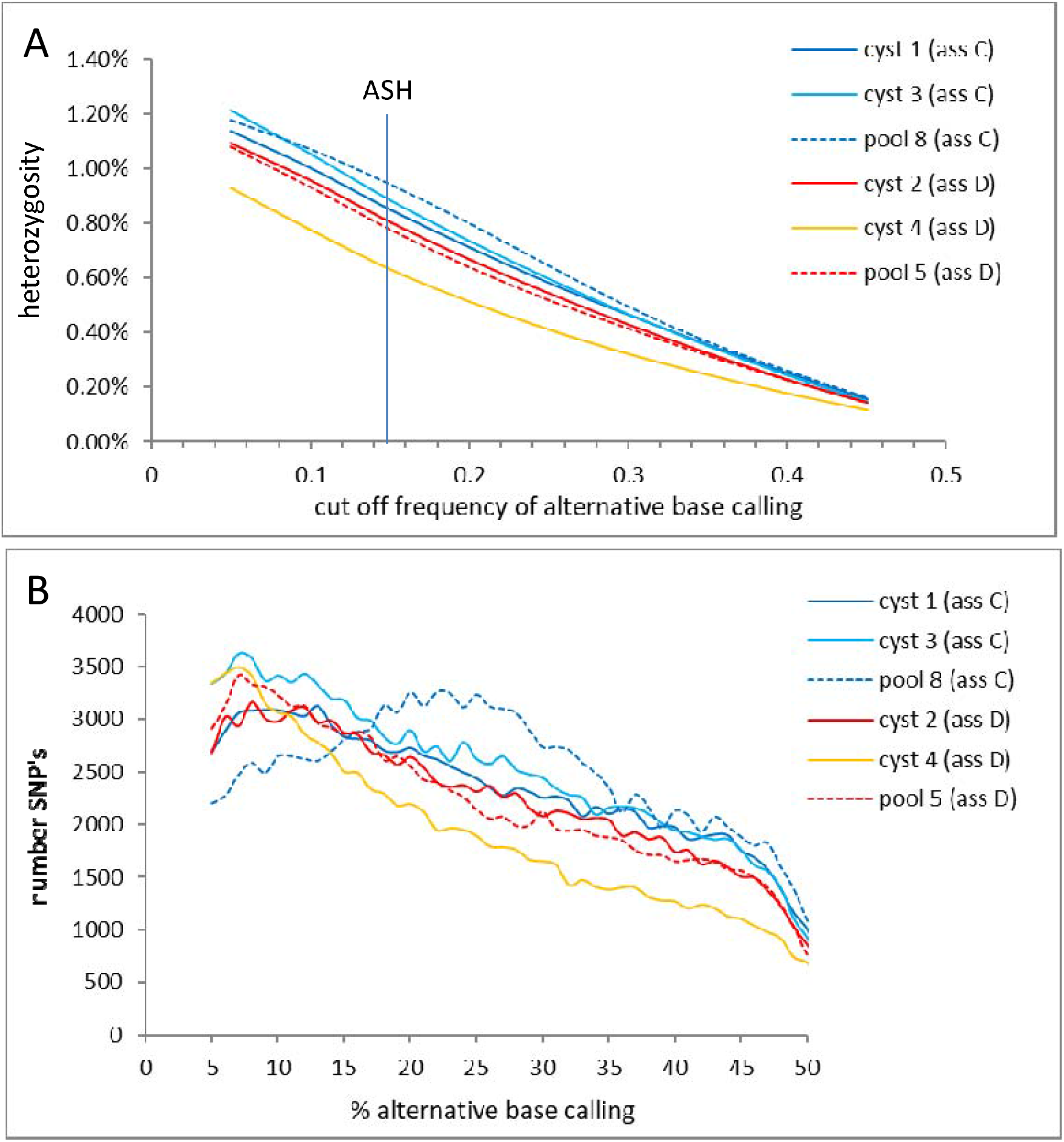
Heterozygosity at different levels of alternative base calling. Heterozygosity as function of the alternative base calling for all 3 genomes of assemblage C (blue lines) and assemblage D (reddish lines) A) Heterozygosity as function of the cut-off frequency of alternative base calling. Heterozygosity at alternative base calling of 0.15 is by definition the allelic Sequence heterozygosity (ASH). B) Number of SNPs for all 3 genomes of assemblage C and D as function of the % alternative base calling.

### Phylogeny

We constructed a phylogenetic tree of concatenated alignments of single copy proteins shared by *G. duodenalis* and *S. salmonicida.* The phylogenetic tree is given in Figure 3. As expected, there is very little difference within the 3 genomes of assemblage C or D. The genomes from assemblages C and D form 1 clade. This in contrast to the genomes of the human assemblages A and B. Assemblage A is more related to assemblage E than to assemblage B. However, the distance between assemblage C and D is comparable to the distance between the assemblages A and B and is larger than between assemblages A and E.

**Fig. 3.**
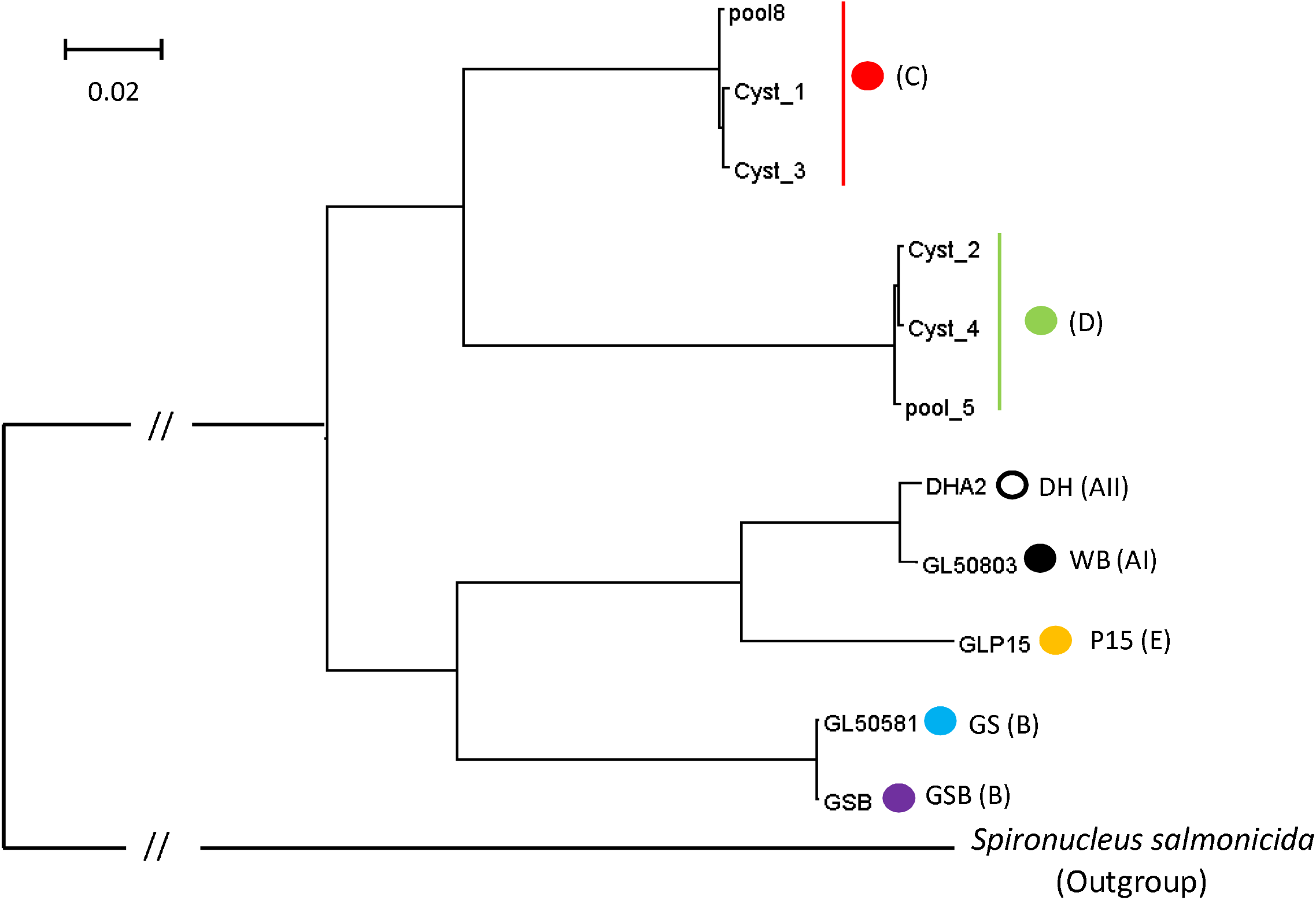
Phylogenetic tree based on genomes from 5 assemblages of G. duodenalis and S. salmonicida. Phylogenetic tree of the *G. duodenalis* genomes of assemblages A (WB and DH), B (GS and GSB), C (cyst 1, cyst 3 and pool 8), D (cyst 2, cyst 4 and pool 5) and E (P15) with *S. salmonicida* as outgroup. Analysis was performed with Maximum Likelihood method. The analysis involved 11 genomes and 534,584 positions. The bar indicates the number of amino acid substitutions per site analyzed. All branches had a bootstrap value of 100.

### Shared and unique OGs of all assemblages

The unique and shared OGs for all assemblages are given in figure 4. For reasons of clarity we used only one genome per assemblage. The GSB genome was preferred over the GS genome, because of the better assembly, resulting in less and longer contigs (Adam et al. 2013). For the same reason the WB genome was preferred over the DH genome (table 1). All assemblages together counted 5508 OGs of which 3288 OGs (59.7%) were shared by all assemblages (core genome). The percentage assemblage specific OGs for assemblage A, B, C, D and E were 14.7%, 9.5%, 10.6%, 9.6% and 1.2%, respectively.

**Fig. 4.**
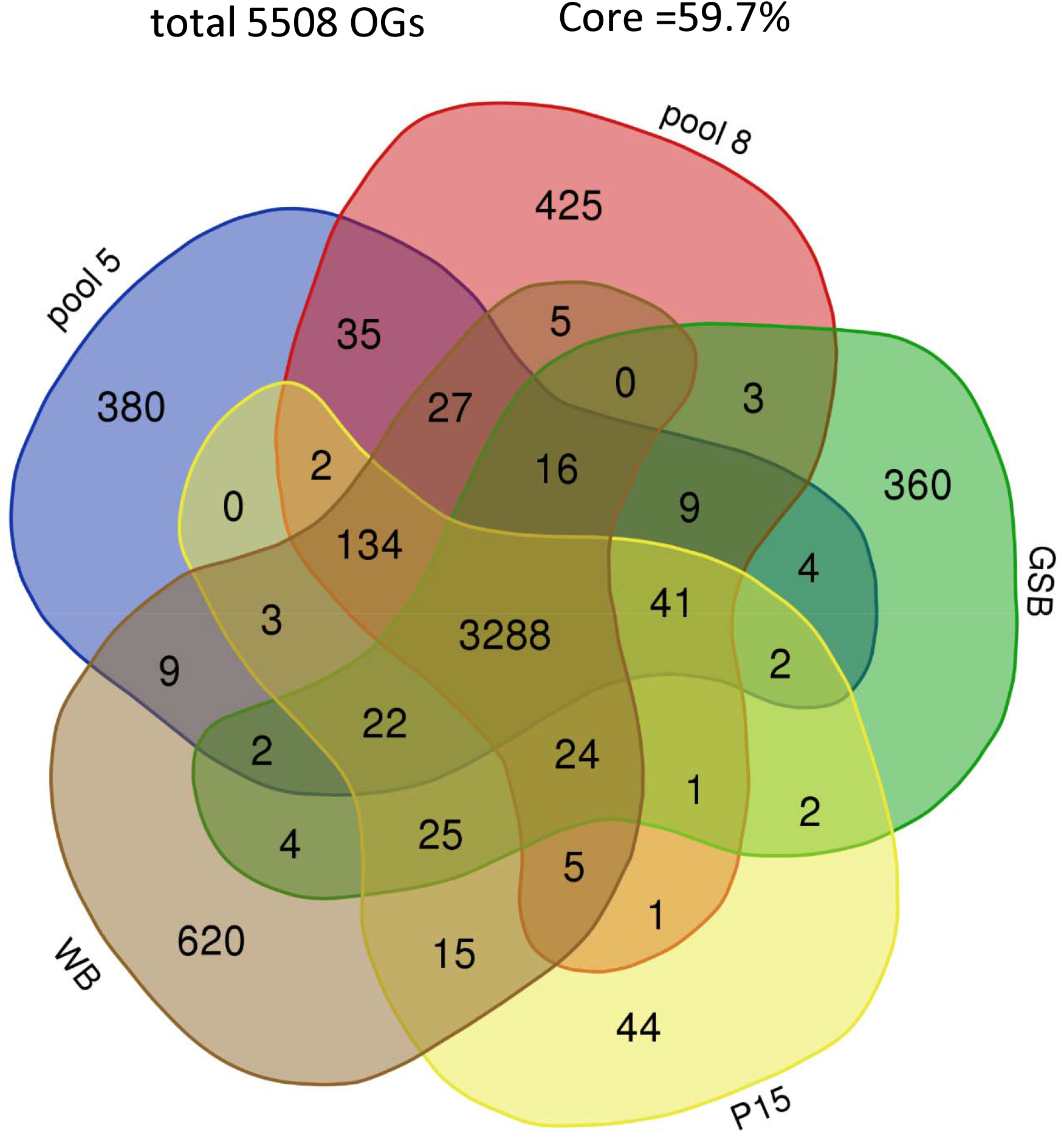
Unique and shared OGs in 5 assemblages. Distribution of the OGs from 5 genomes representing the assemblages AI (WB), B (GSB), C (pool 8), D (pool 5) and E (P15) of *G. duodenalis* shown in a Venn diagram. In total there were 5508 OGs and 3288 OGs (59.7%) were shared between all assemblages.

### OGs unique for dog specific assemblages

OGs present in all the 6 genomes of the dog specific assemblages C and D, but in none of the other genomes are considered dog specific. Initially, 19 OGs were identified as dog specific (Table S2). Manual BLASTp analysis against the non-redundant protein database demonstrated the presence of 5 out of these 19 OGs also in the non-dog assemblages. Four OGs had highest identity with organisms other than *Giardia*. Ten OGs annotated as hypothetical protein did not resulted in a BLAST hit against non-redundant protein database.

### OGs unique for assemblages specific for hosts others than dogs

There were 14 OGs present in all genomes except the genomes of assemblage C and D (Table S3). Two of the genes belonging to this 14 OGs were Flavohemoglobin and a 4Fe-4S binding domain family protein, one of the 5 ferredoxins in *G. duodenalis* (Ansell et al., 2017). Both genes belong to the oxidative response network of Giardia (Ma’ayeh et al., 2015). Flavohemoglobin is upregulated by NO in strain WB (Mastronicola et al., 2010) and by O_2_ in strain WB and GS (Ma’ayeh et al., 2015) and is involved in the detoxification of O_2_ and NO. Ferredoxin mediates electron transfer in the detoxification of O_2_ and NO (Ma’ayeh et al., 2015). The 2Fe-2S, ferredoxin gene (GL50803.27266) was also missed in the automated annotation of the dog assemblages, but that was because this gene contains one of the four introns in the *Giardia* genome. Manual search identified this gene, including the intron, in assemblage C (not shown).

### Cathepsin-B

The cathepsin-like cysteine proteinase family (OG000011) contained in total 139 sequences from all genomes together. Fourteen sequences with a GPI anchor or truncated sequences with high identity to sequences with a GPI anchor were very dissimilar to the other sequences and were therefore removed. From the 125 remaining sequences, those sequences that were not long enough to have a complete peptidase domain (200 amino acids), were removed together with the sequences that lacked the gln-cys-his-asn residues conserved in cathepsin-B (Sajid and McKerrow, 2002). From the remaining 85 sequences a phylogenetic tree was constructed and 9 clades could be identified (Figure 5). Clades 1 to 6 contained 1 gene from each genome, except for clade 5 that missed only the gene from the GS genome. Within these clades the sequence variation between the orthologs was very low. Clades 7, 8 and 9 were more diverse, but closely related to each other. In these clades more than 1 cathepsin gene per genome can be found within one clade, for example 3 cathepsin paralogs of GSB are found in clade 7. On the other hand, not all genomes were represented in all these 3 clades. Clade 8 was the only clade that contained genes from assemblages C and D, but the genes from assemblage A and E were missing in this clade. Therefore, the synteny of cathepsin-B genes of clade 8 was studied by aligning a 47,000 bp fragment with the homologous contigs of the other assemblages (figure 6). Two deletions in the genomes of assemblages A and E were found close to each other, one of 6,300 bp and another of 670 bp, causing the deletion of the chromosome segregation ATPase gene and the cathepsin-B gene, respectively, in the genomes of assemblages A and E.

**Fig. 5.**
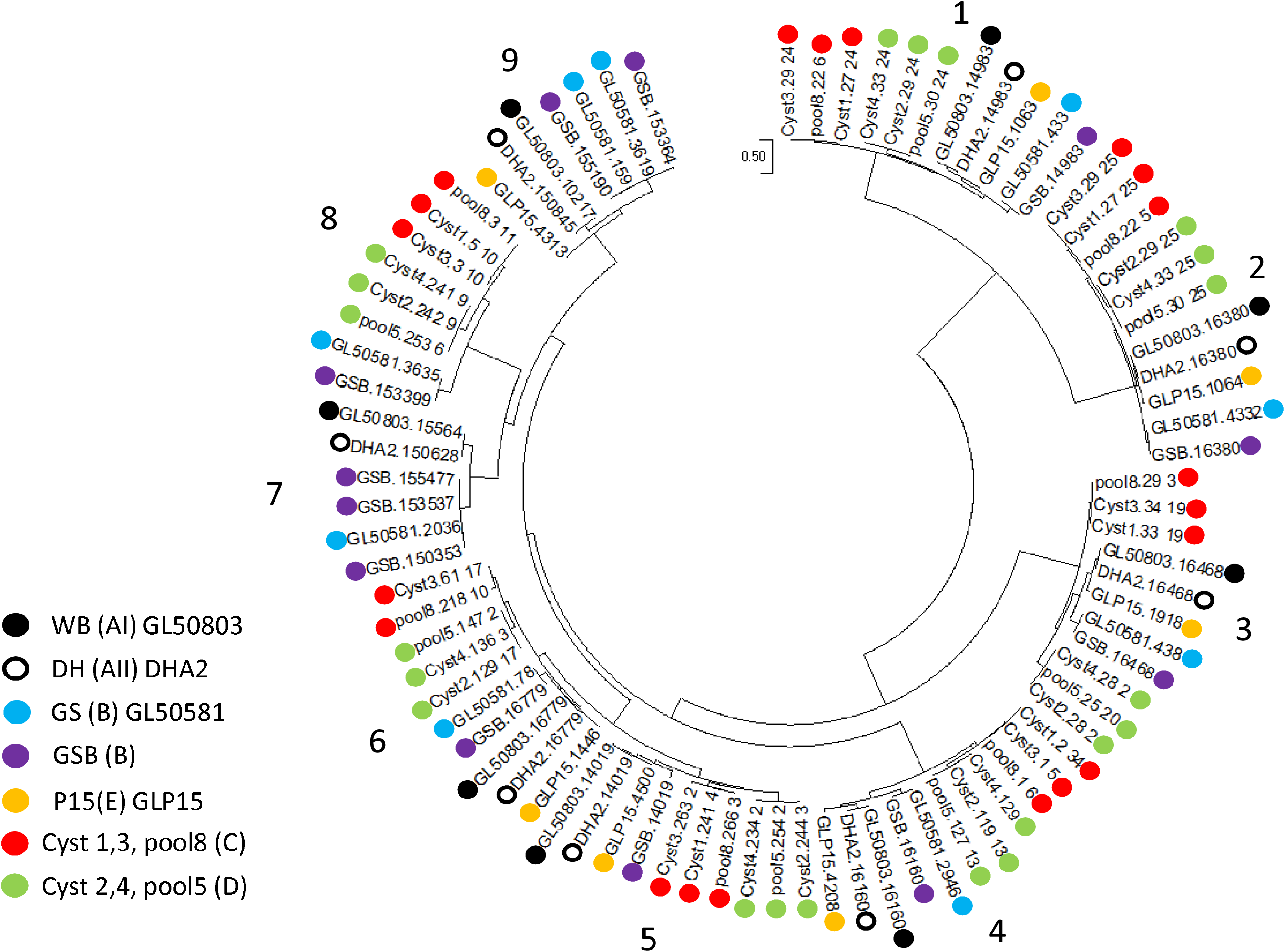
Phylogenetic tree cathepsin-B genes from 5 assemblages. Molecular Phylogenetic analysis by Maximum Likelihood method with gamma distribution of the 85 cathepsin-B genes from 11 genomes, indicated by colored dots; WB (black), DH (open black), GS (blue), GSB (purple), P15 (orange), assemblage C (red) and assemblage D (green). The analysis involved 85 amino acid sequences with in total 620 positions in the final data set. Analysis were conducted in MEGA 6.6.

**Fig. 6.**
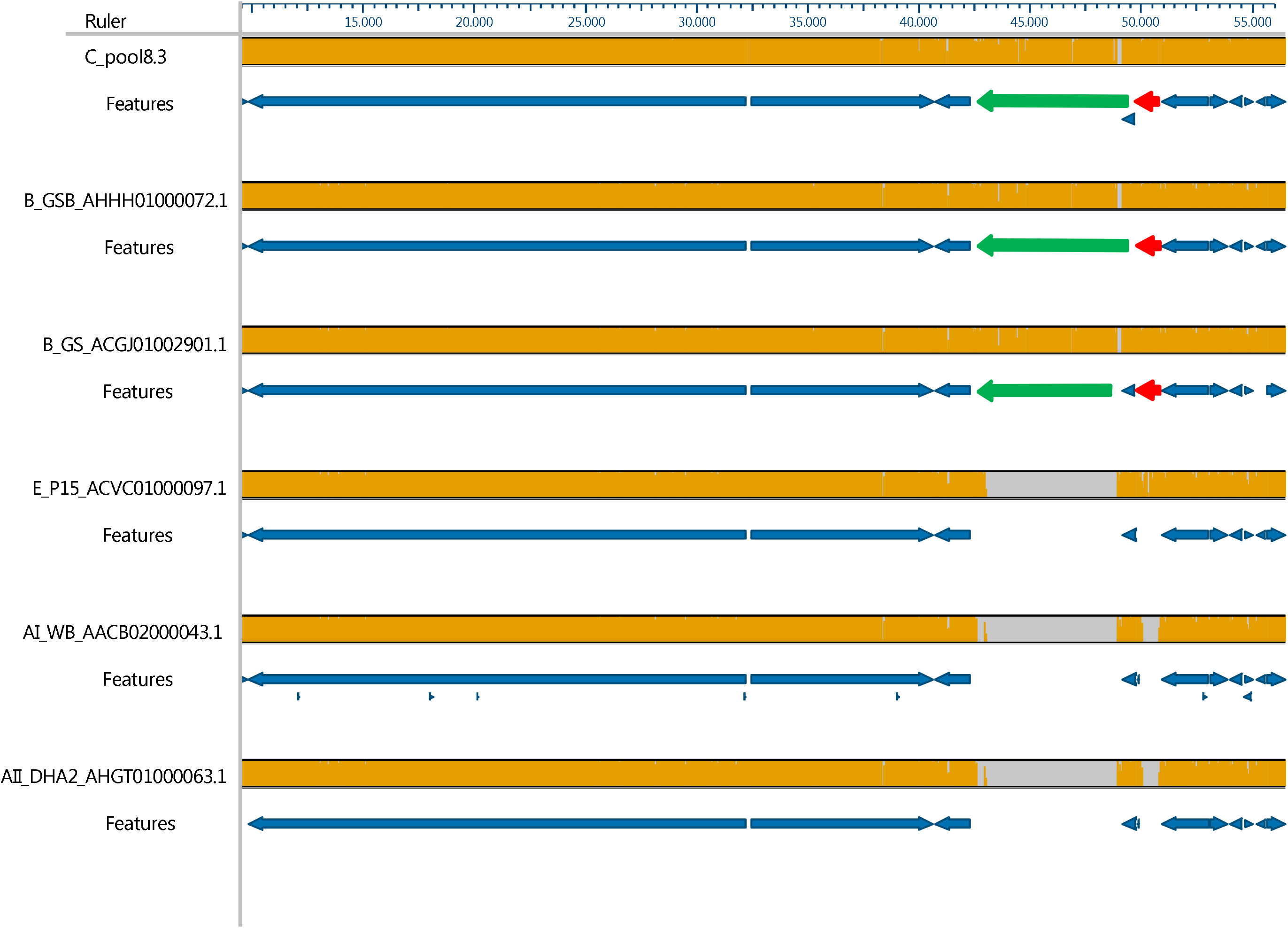
Synteny of the 47,000 bp region around the cathepsin-B gen of clade 8. Synteny study of cathepsin-B genes from clade 8. Homologous contigs of all genomes were aligned in MAUVE. Alignment was restricted to a 47,000 bp fragment. Orange in the bar indicates homology, gray indicates deletions. Features indicate the annotated genes. The chromosome segregation ATPase gene (green arrow) and the Cathepsin-B gene (red arrow) were deleted in the genomes from assemblage E, AI and AII. Pool 5 (assemblage D) was completely homologous with pool 8.3, but is not shown, because the 47,000 bp region consisted out of 3 short contigs, hampering automated annotation.

## Discussion

Whole genome sequences were successfully obtained from the dog specific assemblages C and D of *G. duodenalis*. After an initial step on a sucrose cushion and fixation in ethanol, it was possible to isolate labeled single cysts by FACS. Multiple displacement amplification (MDA) of the genome of single cysts yielded DNA of sufficient quality and quantity to perform whole genome sequencing successfully.

The single cysts of assemblage C and D were isolated from the same dog with a mixed infection. Nevertheless, RFLP performed on the MDA products of these cysts could not demonstrate any contaminating DNA from the other assemblage. The absence of significant contamination was confirmed by the comparison of the genomes obtained from single cysts with the genomes from pooled cysts from dogs with a mono-infection of assemblage C or D. The number of OGs shared between the genomes of individual cysts of different assemblages is not larger than between the genomes of the pooled cysts of the different assemblages. Apart from this, a small number of OGs had to be removed because of bacterial origin, these were all contained on contaminating contigs. This was especially the case in the genome from pool8 of assemblage C. More extensive washing of the cysts and further dilution of the cyst suspension before cell sorting, can likely reduce this contamination even further in next experiments.

The whole genome sequences of assemblage C and D were 99% complete compared to single copy genes from the other genomes of *G. duodenalis.* The completeness of the genomes derived from single cysts was the same as that from 40 pooled cysts. The chance that more than 1 cyst per well were sorted seems unlikely, because of the diluted suspension of cysts and because of the gating, which selects only single cysts. There were on average 115 unique OGs present in the genomes from the pooled cysts and 84 unique OGs in the genomes from single cysts. The higher number of OGs in the pooled samples could be the result of more DNA template being present, but it can also be because the pooled cysts were from a different sample (dog). In any case, the difference was only small. MDA performed on single cells prior to whole genome sequencing was described before for prokaryotes (Ellegaard et al. 2013; Lasken 2007). In order to increase the completeness of the genome, Ellegaard et al. (2013) suggested to pool cells prior to MDA amplification, whereas, Lasken (2007) suggested to pool the MDA products prior to whole genome sequencing. We combined this approach by applying the MDA procedure on 10 pooled cyst, followed by pooling 4 MDA products before sequencing. We found little difference between the genomes obtained from 1 cyst or from 4×10 cysts. Amplification by MDA of individual single celled parasites prior to whole genome sequencing was described before on single oocyst of *Cryptosporidium parvum* (Troell et al. 2016). There, on average 81% of the reference genome could be accounted for in the individual single cell sequence data. Combining 10 individual oocysts before sequencing almost completely covered the whole reference genome. The cysts of Giardia are 16N (Bernander et al. 2001), while the *Cryptosporidium* oocyst are only 4N. This is a possible explanation why the completeness of the genome was in the case of *Giardia* single cell sequencing better than with *Cryptosporidium*, as more DNA is available per reaction.

The genomes of assemblage C and D were of about the same size and contained the same number of genes. 71.7% of the OGs were shared and about 4% were unique for each assemblage. The phylogenetic tree of the genomes demonstrated that assemblage C and D belong to the same clade. This is in agreement with the phylogeny based on a single locus (*gdh*) by Feng and Xiao (2011). This is not the case with the two human assemblages. The genomes from assemblage A are found to be more related to assemblage E, than to the other human assemblage B. This has been found before (Monis et al. 1999). Nevertheless, the divergence between assemblage A and B is similar to that between assemblage C and D and is comparable to that between species of *Leismania* or *Theileria* (Jerlström-Hultqvist et al. 2010b; Adam et al. 2013). This justified the suggestion to change the taxon of assemblage A and B into 2 species, *G. duodenalis* and *G. enterica*, respectively (Thompson and Monis 2012). The same authors suggested to change the dog assemblages C and D into a single new species *G. canis.* By other authors (Ryan and Cacciò 2013) it was suggested not to group the two dog assemblages into one species, but to make 2 species of it because of the differences in sequences of the loci used for multilocus genotyping. The genetic difference between the genomes of assemblage C and D, comparable to the difference between species of *Leismania* or *Theileria,* potentially justifying this division into 2 species. Furthermore, the symptoms seemed to differ between assemblage C and assemblage D infected dogs, with less symptoms in the case of an infection with assemblage C (Pallant et al. 2015). Interestingly, in our study, the dog infected with assemblage C also had less symptoms than the dogs infected with assemblage D or with assemblage C and D.

The allelic sequence heterozygosity (ASH) varies strongly between the assemblages. The lowest ASH is found in assemblage A1 and E, <0.01% and 0.0023%, respectively (Morrison et al. 2007; Adam et al. 2013). The ASH for the assemblages C and D (0.89% and 0.74%, respectively) is slightly higher than for the assemblage B, genome GSB (0.43%) or GS (0.53%) (Adam et al. 2013; Franzén et al. 2009). Although it is difficult to compare the ASH of the different genomes, because different sequencing methods are used, the difference in ASH between A and E on one side and B, C and D on the other side is striking. The same division of the assemblages can be seen in the phylogenetic tree with assemblage E and A forming one clade. It suggests that having a low ASH is a derived character from a predecessor with a high ASH as found in assemblage B, C and D. The ASH of assemblage F, G and H are not known. Given the position of assemblage F in between assemblage A and E in the phylogenetic tree (Feng and Xiao 2011), it is predicted that it will have also a low ASH. Likewise, assemblage G and H located on the base of the rooted phylogenetic tree, is predicted to have a high ASH. In the present study, a linear decrease in heterozygosity was seen as function of the cut-off frequency of alternative base calling. If a 16N cyst consists of nearly identical haploid genomes with random SNPs this could be expected and this was actually found in 5 out of the 6 genomes. Only the genome obtained from de pooled cysts of pool 8 (assemblage C) showed a peak in the number of SNPs at the 25% alternative base calling. This suggests the presence of 4 haploid genomes with equal abundancy, like the 4 haploid genomes present in a trophozoite. Why this is seen only in the genome of pool 8 and not in the other genomes, is difficult to explain, however it could be that much higher sequencing depths are needed to distinguish alternative base calling frequencies.

After manual BLASTp search, 14 OGs were identified as dog specific. Ten of these OGs coded for a hypothetical proteins for which no hits were found in GenBank. For four OGs the hits with the highest E value were bacterial genes coding for annotated proteins. Further research is needed if these genes play a role in host adaptation.

There were 14 OGs found in all genomes of assemblage A, B and E, but lacking in the dog assemblages C and D. Two genes of this group are flavohemoglobin and 4Fe-4S binding domain family protein, a ferredoxin. Both proteins are part of the oxidative response network (Ma’ayeh et al., 2015). Flavohemoglobin and ferredoxins play a role in detoxification of O_2_ and NO. Flavoprotein (OG000298) and other ferredoxins were present in all genomes and this makes possibly flavohemoglobin and 4Fe-4S binding domain family protein redundant in the dog assemblages C and D.

Cathepsins-B from *Giardia* are known to play a role in host-parasite interaction. Several orthologs have been demonstrated to be excreted (Dubourg et al. 2018) and digestion by cathepsins-B of host IL-8 (Cotton et al. 2014) and of proteins involved in tight junction of host epithelial cells and several chemokines (Liu et al. 2018) have been described. The phylogenetic tree of the cathepsin-B sequences from all 11 *Giardia* genomes consisted of 9 clades. Six clades showed little variation between the orthologs, suggesting that these genes have important functions in all genomes, independent of the assemblage, and therefore independent of the host. Three Clades (7, 8 and 9) showed a larger diversity. Nothing was known so far about the cathepsins from the dog assemblages C and D. We found seven cathepsin-B genes in both dog assemblages and the orthologs from assemblage C and D were very similar to each other. The cathepsin-B genes from assemblage C and D were present in the conserved clades 1 to 6 and in the variable clade 8, but were missing in clades 7 and 9. Synteny study demonstrated the deletion of the orthologs of the clade 8 cathepsins in the genomes of the assemblages A and E. The large variation within clades 7, 8 and 9 can indicate that these cathepsins do not have important functions, but it can also be the result of a different function in the different assemblages infecting different hosts. A recent finding suggests that the cathepsins from the variable clades indeed have an important function. It was shown that the orthologs from clade 7, GL50803.15564 and GL50581.2036 from genome WB and GS, respectively, have dN/dS values of >26, indicating very strong positive selection (Dubourg et al. 2018). This strong selection is not expected when these genes do not have an important function. The formation of the 6 paralogs of genome GSB within clades 7, 8 and 9 can also be caused by selection, because it is known that substrate specificity varies between the paralogs and orthologs of the cathepsins of *Giardia*. Recombinants of 3 paralogs of cathepsins from GL50803 (Liu et al. 2018), but also orthologs of GL50803 and GL50581 (Bhargava et al. 2015, Cotton et al. 2014) showed differences in substrate specificity. Therefore, the presence of more cathepsin variants within an assemblage, might help to make a less restricted host repertoire possible. The absence of genes from assemblage C and D in clade 7 and 9 might therefore be related to this more restricted host repertoire of these assemblages, compared to the 2 to 6 genes of the assemblage A and B with a broader host range (Sprong et al. 2009). It suggests that study of the substrate specificity and other characteristics of the cathepsins-B of clade 7, 8 and 9 and other cathepsins-B can reveal inside into some aspects of the host specificity.

## Acknowledgements

We are grateful to the Veterinary Microbiological Diagnostic Centre (VMDC, Faculty of Veterinary Medicine, Utrecht, University, the Netherlands) for the samples of Giardia positive dogs. Kristel van Rooijen is thanked for her technical assistance by the isolation of the individual cysts. Dr Ger Arkesteijn is thanked for the FACS analysis.

